# An anti-SARS-CoV-2 formula consisting volatile oil from TCM through inhibiting multiple targets in cultured cells

**DOI:** 10.1101/2024.05.22.595089

**Authors:** Huan Zhang, Wuyan Guo, Jing Shen, Xiaorui Song, Bo Zhang

## Abstract

At present, coronavirus severe acute respiratory syndrome coronavirus 2 (SARS-CoV-2) and its variant virus are prevalent all over the world, taking a major toll on human lives worldwide. Oral mucosa and saliva are the high-risk routes of transmission. Inactivation of the virus in the mouth is one of the important strategies to reduce the source infectivity of the virus. However, the current preparations for inactivating oral virus are mainly aimed at ACE2 protein, and its effect needs to be improved. In this study, through literature mining, multi-target screening and other methods, we screened and optimized the volatile oil formula (*Artemisiae argyi folium, Chrysanthemum morifolium, Trollii chinensis flos, Lonicerae japonicae flos*) from the volatile oil of traditional Chinese medicine to inhibit the invasion and replication of SARS-CoV-2. A test using pseudotype virus with S glycoprotein confirmed that this formula could effectively bind to the S glycoprotein to prevent SARS-CoV-2 host cell entry. Molecular docking experiments showed that the multiple molecules in the formula might weakly bind to these targets including ACE2, cathepsin L, furin, mpro/3cl. In all, the volatile oil formula designed in this paper offers a general affordable strategy to protect patients from SARS-CoV-2 infections through debulking or minimizing transmission to others.

## Introduction

Globally, the severe acute respiratory syndrome coronavirus 2 (SARS-CoV-2) outbreak has caused millions of infections and tens of thousands of deaths. The spread of SARS-CoV-2 occurs both through droplets and aerosols, and is strongly linked to indoor exposures of infected individuals, whether they are symptomatic or not^[1]^. In saliva, high levels of SARS-CoV-2 virus are often detected^[2]^. Both asymptomatic and symptomatic COVID-19 patients have high SARS-CoV-2 viral loads in saliva^[3,4]^. In fact, COVID-19 symptoms include the loss of taste and smell, and viral replication occurs in salivary glands and oral mucous membranes^[5]^. Therefore, Virus inactivation within the oral cavity may be an effective strategy to reduce the spread of SARS-CoV-2 through the oral mucous membranes and saliva.

A better understanding of how SARS-CoV-2 hijacks the host during infection is essential for constructing therapeutic strategies. In fact, SARS-CoV-2 replication relies on many proteins or enzymes from the virus itself and the host cell^[6]^. In principle, these proteins and enzymes represent potential therapeutic targets. Coronaviruses start their life cycles when their S proteins bind to angiotensin-converting enzyme 2 (ACE2) on host cells’ surface membranes^[7]^. According to a previous description, the pre-fusion conformation is initiated by the proteolytic cleavage of the S1/S2 site, catalyzed by serine-proteases like furin^[8]^ or cathepsin L^[9]^ in correspondence of the (R/K)-(2X)n-(R/K) motif of S. As a result of furin cleavage of the S1 (SPRRARY S) site of SARS-CoV-2, spike glycoprotein activation takes place, allowing the virus to enter the host. It is a polybasic cleavage site found in SARS-CoV-2, also known as a multibasic cleavage site. In COVID-19 patients, Cathepsin-L is thought to play a critical role since its levels are elevated after SARS-CoV-2 infection and positively correlate with disease severity^[10]^. Functionally, SARS-CoV-2 infection is enhanced by cathepsin-L by cleaving furin-primed SARS-CoV-2 S protein into smaller fragments and promoting cell-cell fusion^[10,11]^. Subsequently, the coronavirus main proteinase (3CLpro) of the virus cleaves viral replicase polyprotein into effector proteins^[12]^. Considering their undeniable role in the entry and replication of viruses, it may be possible to inhibit or modulate these four proteins (ACE2, furin, cathepsin L and 3CLpro) to develop effective antiviral therapies.

Currently, the oral preparation used to inactive SARS-CoV-2 are mainly aiming to act on the ACE2 target, which plays a major role in the invasion of host. Daniell et al developed a chewing gum with converting enzyme 2 proteins to decrease oral virus transmission and infection through debulking SARS-CoV-2 in saliva^[13]^. Another study demonstrated newly synthesized polypeptides inhibits eliminating the invading virus by competing with SARS-CoV-2 for the ACE2 receptors^[14]^. However, these Oral preparation have been exclusively focused on one single protein target. Based on the fact that viruses go through various stages of their life cycle, it is ideally to target multiple viral proteins in that it enables the potential preparation to disrupt the viral infection and replication process in different stages^[15]^, as a result, the effect of killing viruses is maximized.

Recent studies showed that Traditional Chinese Medicine (TCM) could acts on multiple targets and multiple pathological processes^[16]^, which provided us a valuable resource. Traditional Chinese medicine volatile oil rich in ketones, terpenes, aldehydes, alcohols and other active substances is the product of plant metabolism extracted from plants by specific methods^[17]^. It is widely used in development of drugs, antibacterial agents, antiviral agents and other products due to its activities of enhancing immunity, bacteriostasis and antiviral^[18,19]^. Moreover, it has aromatic components, and the raw materials are simple and easy to obtain. Compared with traditional chemical drug products, volatile oil can possess more efficient antibacterial activity and are not easy to produce resistance limitation, with lower cytotoxicity^[19,20]^. Thus, it is feasible to develop safe and environment-friendly anti-SARS-COVID-2 oral preparation from the volatile oil of traditional Chinese medicine.

In this paper, we screened from the volatile oils of traditional Chinese medicine based on literature mining and multiple target screening and obtain an effective formula. Then, the effect of the formula was investigated by *in vitro* assay. Finally, we use molecular docking to preliminarily explore the mechanism of this compound against SARS-CoV-2.

## Materials and methods

### Reagents and Cell Culture

HEK293T□overexpressing ACE2 cell were purchased from GENEWIZ (Jiangsu, China) and grown in DMEM medium (Gibco, Thermo Fisher Scientific, Waltham, MA, USA) containing 10% fetal bovine serum (Gibco; hereinafter referred to as complete medium), 100 U/mL penicillin, 100 μg/mL streptomycin and 1 μg/mL puromycin and incubated at 37°C in an atmosphere containing 5% CO2/95% air.

### Literature Mining

“Traditional Chinese medicine”, “volatile oil” and “antiviral” were used as key words for a comprehensive search of both English- and Chinese-language literature in PubMed, and CNKI (Chinese National Knowledge Infrastructure) databases from 2000 to May 2022. For overlapping or republished studies, only the most recent published papers were included. There are 214 references and 62 kinds of single traditional Chinese medicine volatile oils are retrieved. Finally, 9 kinds of traditional Chinese medicine volatile oils was obtained according to the *in vitro* antiviral mechanism.

### Extraction of Volatile Oil by Hydrodistillation

Five kg of 9 kinds TCM (Yu xing cao, Jin lian hua, Hong feng ye, Jin yin hua, Shi xiang ru, Ma ye qian li guang, Hang bai ju, Ye ju hua, Ai ye) were purchased from Anguo Kanghua Traditional Chinese Medicine sales Co. Ltd. (Hebei, China).

Each of above TCM was subjected to hydrodistillation for 2 h, to extract volatile oil, and the volatile oil was then kept in a glass bottle after cooling down to the room temperature. The residual water was then removed, using anhydrous sodium sulfate. The yield of essential oil was calculated by using the following equation: %Yield = (a/b) × 100, where a is a volume of the volatile oil, and b is the weight of fresh plant materials used in the hydrodistillation. The essential oil was stored at 4°C, in a light-protected container, until further use.

### Cathepsin L Activity Assays

The Cathepsin L Activity Screening Assay Kit (Biovision, California, USA) was used to determine the inhibitory effect of selected volatile oils on the activity of isolated cathepsin L enzyme according to the manufacturer’s protocol. Selected volatile oils at 0.5 mg/mL concentrations were added to cathepsin L (0.2 mU/μl) and the reaction mix was incubated for 15 min at RT. Positive control consisted of cathepsin L alone, whereas negative control consisted of cathepsin L and inhibitor (10 μM) of cathepsin L. Cathepsin L substrate was added to each well. The fluorescence was measured at Ex/Em = 400/505 nm wavelength using a spectrofluorometer (thermofisher, bMassachusetts, USA) in a kinetic mode for 30 min at 37°C. Control was 0.5% ethanol. Results are expressed as a percentage of volatile oil-free control (mean +/- SD, n = 6).

### ACE2 Activity Assays

To determine the inhibitory effect of selected volatile oils on the activity of recombinant hACE2 protein, an ACE2 Activity Screening Assay Kit (Beyotime, Shanghai, China) was used according to the manufacturer’s protocol. Briefly, the volatile oils at 0.5 mg/mL concentrations were added and the reaction mix was incubated for 45 min at RT. The positive control was a sample containing ACE2 enzyme and ACE2 enzyme inhibitor MLN-4760 (10 nm). The fluorescence was measured at Ex/Em = 325/393 nm wavelength using a spectrofluorometer (thermofisher,bMassachusetts, USA). 100% enzyme activity control was 0.5% ethanol. Results are expressed as a percentage of volatile oil-free control (mean +/- SD, n = 6).

### Furin Activity Assays

To examine the effect of various selected volatile oils on furin activity, a furin Activity Screening Assay Kit (Biovision, California, USA) was used according to the manufacturer’s protocol. Briefly, the volatile oils at 0.5 mg/mL concentrations were added. furin substrate was added to each well. The positive control was a sample containing furin enzyme and furin inhibitor (10 nm). The fluorescence was measured at Ex/Em = 360/460 nm wavelength using a spectrofluorometer (thermofisher,bMassachusetts, USA) in a kinetic mode for 1 h at 37°C. Results are expressed as a percentage of volatile oil-free control (mean +/- SD, n = 6).

### Mpro/3CL Activity Assays

To determine the inhibitory effect of selected volatile oils on the activity of Mpro/3CL protein, an Mpro/3CL Activity Screening Assay Kit (Beyotime, Shanghai, China) was used according to the manufacturer’s protocol. Briefly, the volatile oils at 0.5 mg/mL concentrations were added and the reaction mix was incubated for 45 min at RT. The positive control was a sample containing Mpro/3CL enzyme and Mpro/3CL enzyme inhibitor Ebselen (2 μm). The fluorescence was measured at Ex/Em = 325/393 nm wavelength using a spectrofluorometer (thermofisher,bMassachusetts, USA). 100% enzyme activity control was 0.5% ethanol. Results are expressed as a percentage of volatile oil-free control (mean +/- SD, n = 6).

### Viability Assay

In order to assess cell viability, we used the CCK-8 assay. Briefly, HEK-293T/ACE2 cells were seeded into a 96-well plate at a cell density of 6,000 per well, and allowed to adhere for 24h, followed by treatment with different concentrations of selected volatile oils for up to 48h. Afterwards, complete growth medium was replaced with a fresh one substituted with 10 μl CCK-8, followed by 3 hours at 37 °C incubation. The absorbance was measured at 450 nm using a spectrophotometer (Molecular Devices, San Jose, CA). Results are expressed as a percentage of volatile oils-free control (mean +/- SD, n = 6).

### Pseudovirus Preparation and Neutralization Assay

For pseudovirus neutralization assay, 100 TCID50 per well pseudovirus in medium without FBS was incubated with volatile oils combination for 30 min at room temperature. Then the mixtures (100 μL) were used to infect HEK-293T/ACE2 cells rinsed with PBS. After 5 h, 100 μL medium with 5% FBS was added. After incubation for another 48 h, After 48 h, cells were lysed using Firefly Luciferase Assay Kit (Genescript, catalog no. L00877C), and relative light units were measured using a 96 Thermo Microplate luminometer. Percent neutralization was calculated using GraphPad Prism 8.

### Molecular Docking

Ligand preparation: for molecular docking studies, The structures of the all ligands from table 2-5 were retrieved from EMBL-EBI (www.ebi.ac.uk/chebi/advancedSearchFT.do), in a three-dimensional structure file (SDF) format, and furthermore, the structure was refined using the LigPrep module in Schrodinger’s Maestro (v 12.8). The OPLS4 force field was applied, and 32 different states of stereoisomerism were derived (Schrödinge Release 2021-2: LigPrep, Schrödinge, LLC, New York, NY, 2021).

Protein preparation: we need to evaluate the furin (5JXH), Mpro (6W63), Cathepsin L (3OF9) and antiACE2 (1R42) activity against different ingredients in the each volatile oils computationally, respectively. The three-dimensional structure of proteins was retrieved from the database of Protein Data Bank (PDB). The X-ray crystallographic structures were imported into Maestro using the protein preparation wizard, and this module helps to solve the missing hydrogen bonds, create the disulfide bonds, and optimize (Schrödinge Release 2021-2: Protein Preparation Wizard; Epik, Schrödinge, LLC, New York, NY, 2021; Impact, Schrödinge, LLC, New York, NY; Prime, Schrödinge, LLC, New York, NY, 2021).

The molecular docking was performed using the Glide package (ligand docking) in the Schrodinger suite. The standard precision docking method was applied and performed postdocking minimization and analyzed the results in pose viewer Schrödinge Release 2021-2: Glide, Schrödinge, LLC, New York, NY, 2021

### Statistical Analysis

All data in experiments were analyzed using Prism 8.0 software and are presented as mean ± SEM. Statistical significance was assessed by one-way analysis of variance (ANOVA). *P* value<0.05 indicates significance.

## Results and Discussion

### Validation of TCMs Volatile Oil for Antiviral Activity

Antiviral research has been an active field in recent decades. To date, At present, many traditional Chinese medicines have been reported to have antiviral effects. In the long history of Chinese medicine, some TCMs volatile oil have been reported to have antiviral benefits. Thus, TCMs volatile oil are great candidates for screening and verifying their antiviral properties. To choose TCM volatile oil candidates for validation, we searched the National pubmed as well as China National Knowledge Infrastructure (CNKI) for antiviral TCMs volatile oil. The mechanism of traditional Chinese medicine can be divided into direct inhibition and indirect inhibition. The direct inhibition is mainly to block a certain link in the process of virus reproduction to defeat pathogen propagation; Indirect inhibition is to stimulate and mobilize the immune defense system of the body to play an antiviral role ^[21]^. Since we only consider the direct antiviral effect, that is, to inhibit the virus invasion and replication in the host, we finally get 9 Chinese herbal volatile oils through screening, as shown in Table 1.

**Table 1.**
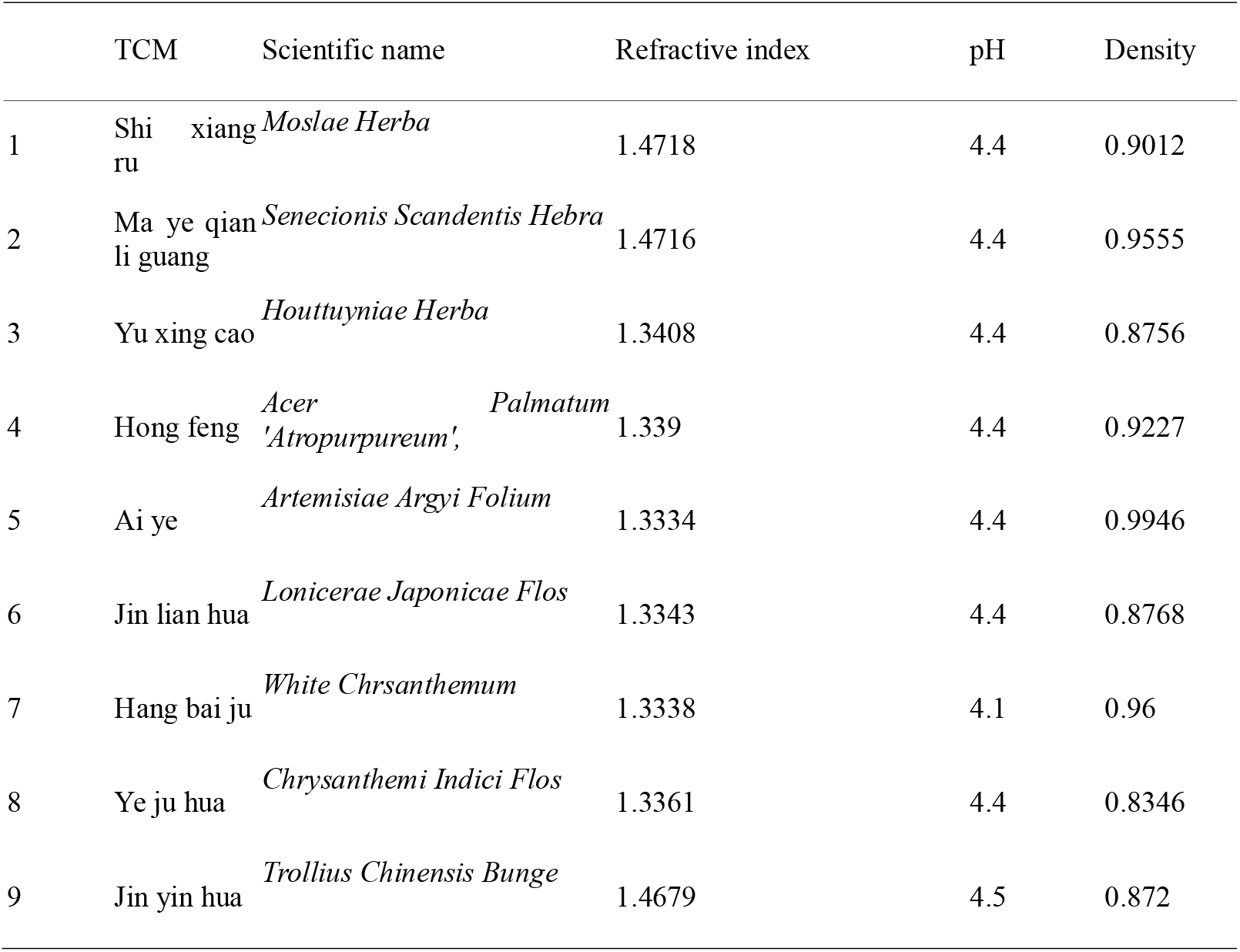
Physicochemical characteristics of 9 volatile oils.

|  | TCM | Scientific name | Refractive index | pH | Density |
| --- | --- | --- | --- | --- | --- |
| 1 | Shi xiang ru | *Moslae Herba* | 1.4718 | 4.4 | 0.9012 |
| 2 | Ma ye qian li guang | *Senecionis Scandentis Hebra* | 1.4716 | 4.4 | 0.9555 |
| 3 | Yu xing cao | *Houttuyniae Herba* | 1.3408 | 4.4 | 0.8756 |
| 4 | Hong feng | *Acer Palmatum*<br>'Atropurpureum', | 1.339 | 4.4 | 0.9227 |
| 5 | Ai ye | *Artemisiae Argyi Folium* | 1.3334 | 4.4 | 0.9946 |
| 6 | Jin lian hua | *Lonicerae Japonicae Flos* | 1.3343 | 4.4 | 0.8768 |
| 7 | Hang bai ju | *White Chrsanthemum* | 1.3338 | 4.1 | 0.96 |
| 8 | Ye ju hua | *Chrysanthemi Indici Flos* | 1.3361 | 4.4 | 0.8346 |
| 9 | Jin yin hua | *Trollius Chinensis Bunge* | 1.4679 | 4.5 | 0.872 |

### Four Volatile Oils Inhibit the Activity of Different Targets of SARS-COV-2, Respectively

As the life cycle of SARS-CoV-2 is related to multiple protein targets, as shown in Figure 1, in order to better prevent SARS-CoV-2 from invading and replicating, we screened volatile oils that can inhibit these four targets, including ACE2, furin, cathepsin L and Mpro/3CL. The angiotensin-converting enzyme II (ACE2) has been recognized as the host receptor of severe acute respiratory syndrome coronavirus 2 (SARS-CoV-2). There is a strong interaction between the receptor binding domain of the SARS-CoV-2 S-protein and ACE2, in this way, ACE2 may provide an effective drug target for preventing COVID-19 from invading host cells[22]. The screening results are shown in Figure 3A. The volatile oil of Hang bai ju is the best one to target ACE2.

**Fig 1.**
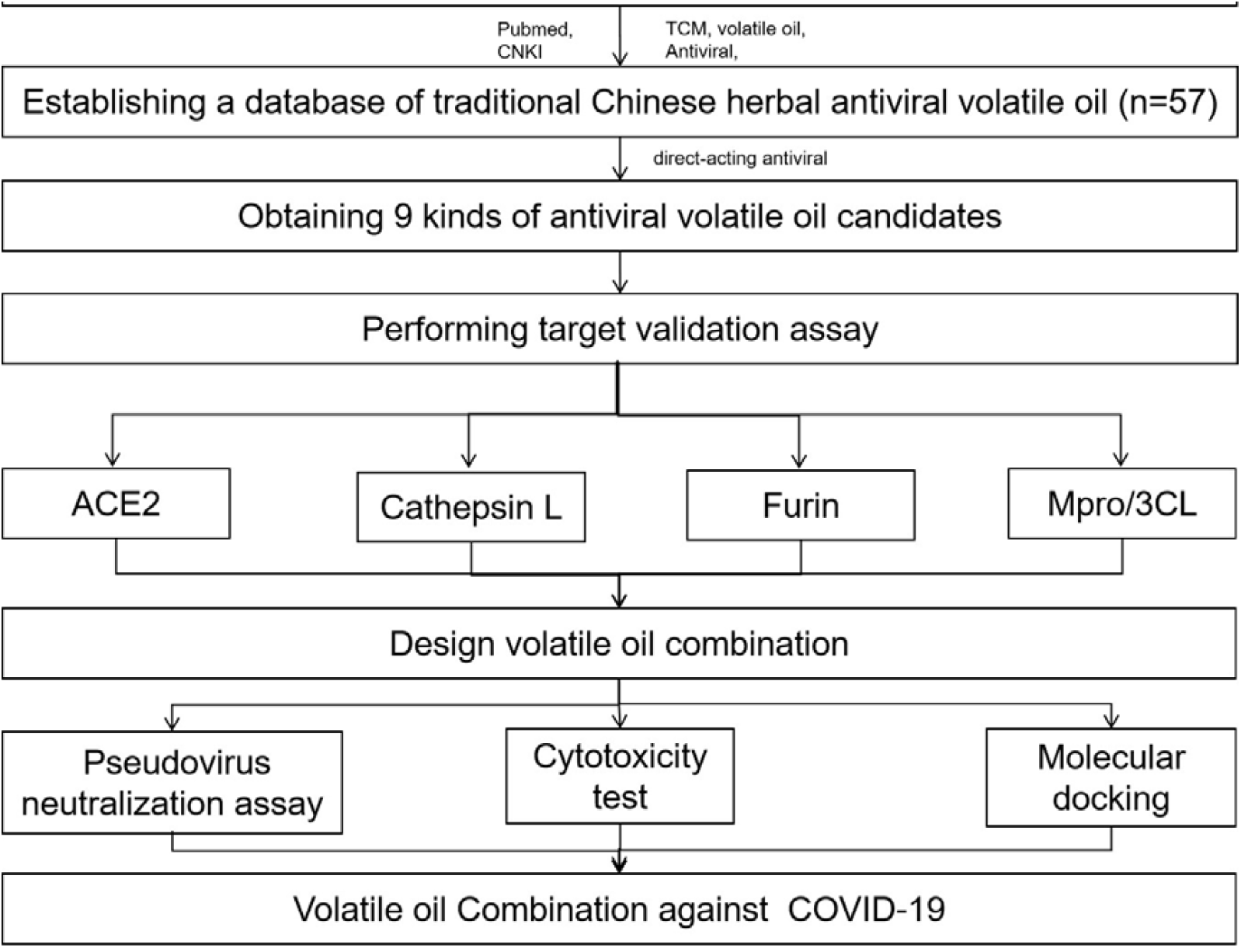
Workflow of this study.

**Fig 2.**
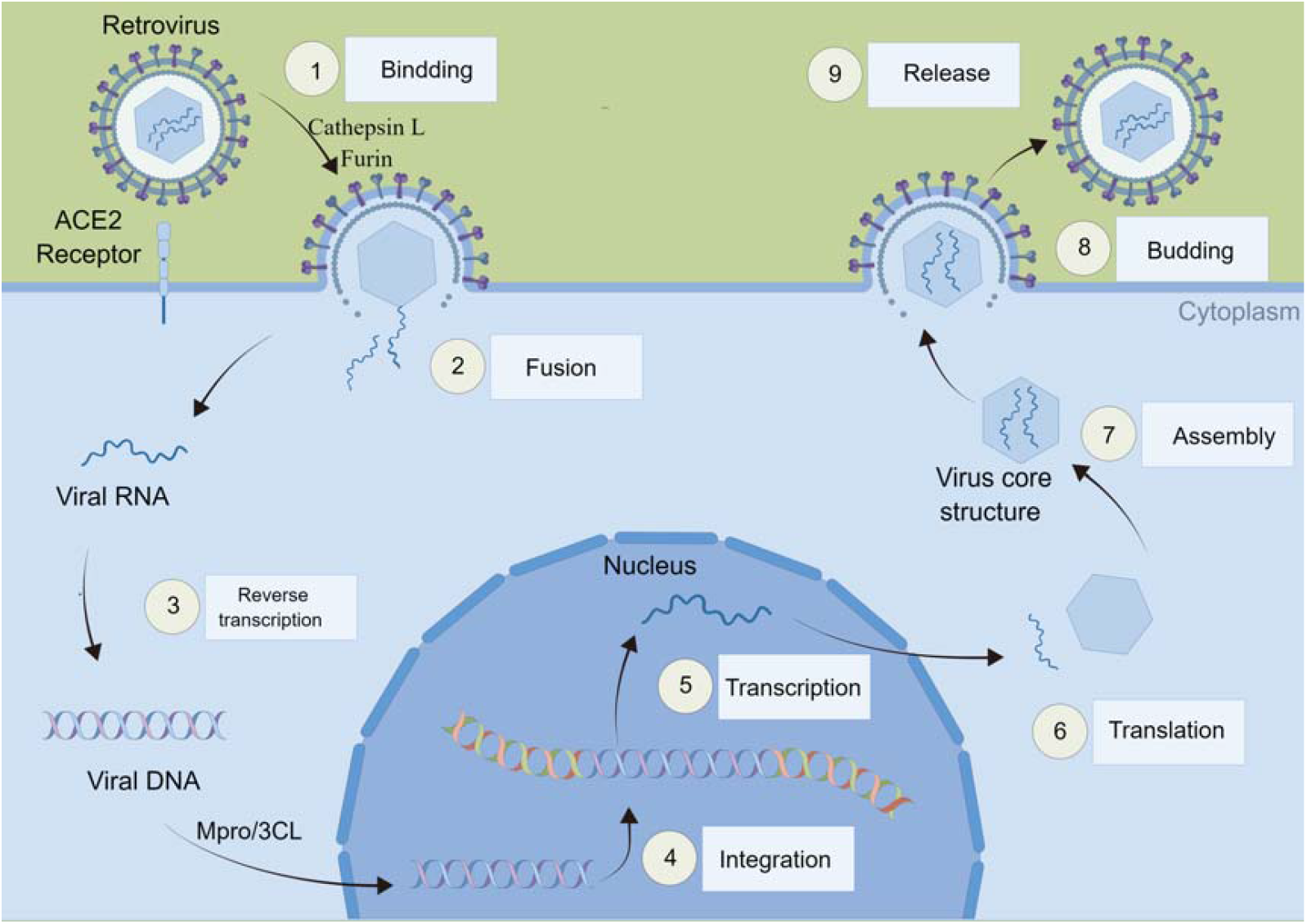
SARS-CoV-2 infection cycle.

**Fig 3.**
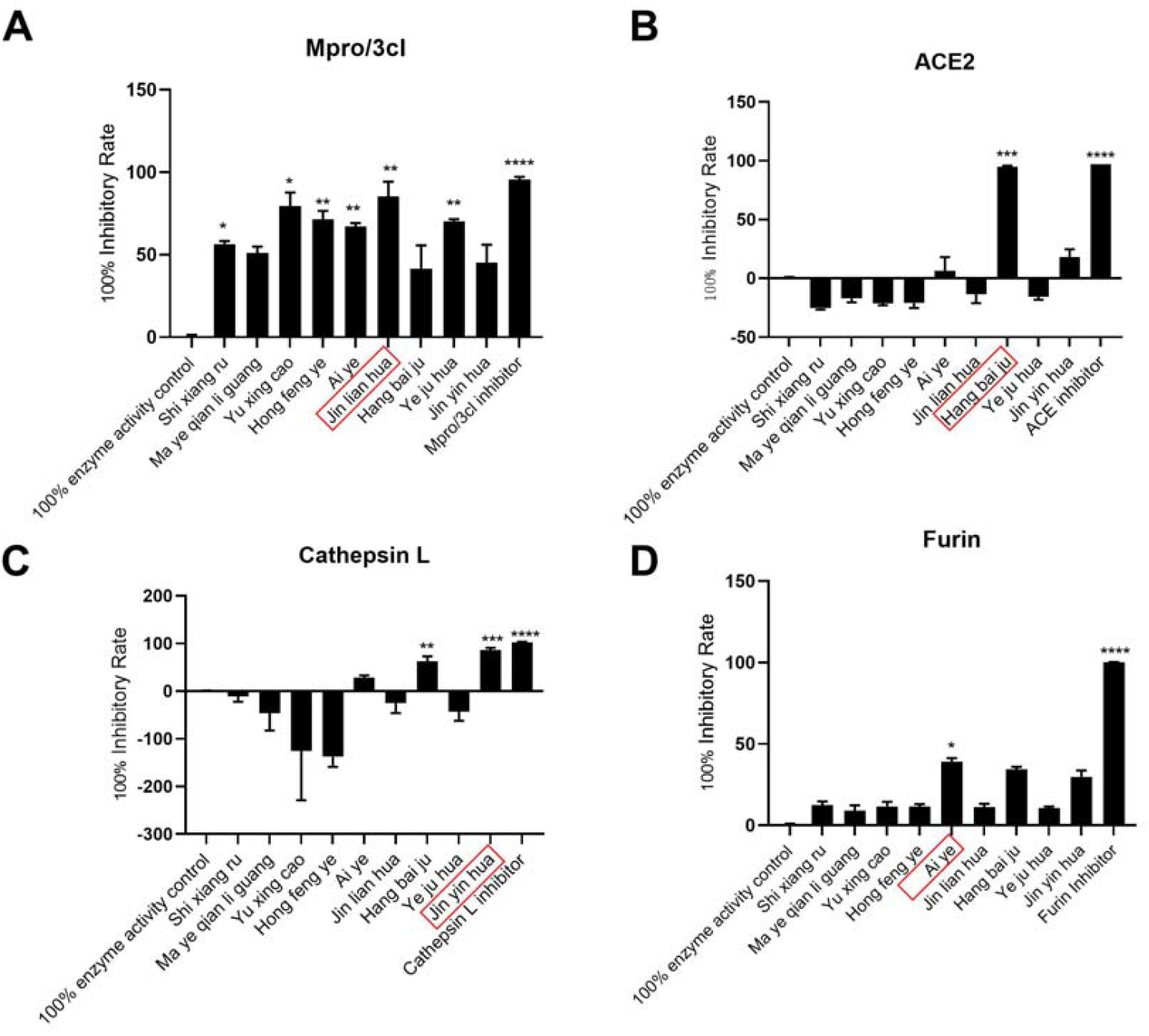
Target screening of volatile oil against SARS-CoV-2. (A) Jin lian hua inhibited the activity of Mpro/3cL at maximal levels. (B) Hang bai ju inhibited the activity of ACE2 at maximal levels(C) Jin yin hua inhibited the activity of Cathepsin L at maximal levels. (D) Ai ye inhibited the activity of Furin at maximal levels. The data represent results combined from three independent experiments. * p < 0.05, ** p < 0.01, *** p < 0.001 compared to 100% enzyme acitvity control.

Furin converts synthesized proteins into biologically active forms by cleaving a specific section[23]. It is a serine endoprotease that cleaves precursor protein processing sites in a calcium-dependent manner. Furin cleaves S1 (SPRRARY S) site of SARS-CoV-2, direct emergence in spike glycoprotein activation for viral entry into the host system[24]. The screening results are shown in Figure 3B. Among the volatile oils of traditional Chinese medicine targeting furin protease, the volatile oil of Ai ye has the best inhibitory effect.

Cathepsin-L plays a critical role in COVID-19 patients, as evidenced by high circulating levels after SARS-CoV-2 infections, and a positive correlation with disease severity[25]. Functionally, It helps promote SARS-CoV-2 infection by fragmenting furin-primed SARS-CoV-2 S proteins and facilitating cell-to-cell fusion by cleaving them into smaller fragments[9,11]. Results as shown in Figure 3C, the inhibitory effect of honeysuckle volatile oil on cathepsin L was the best.

Mpro is highly conserved among different CoVs[26], and as far as we known, It has no human homolog, and shows a low mutation rate, making it ideal for the development of multi-viral drugs that can interfere with different varieties of SARS-CoV-2 and other coronaviruses’ vital cycles[27]. The screening results are shown in Figure 3D. Among the volatile oils of traditional Chinese medicine targeting the main protease, the volatile oil of Jin lian hua has the best inhibitory effect.

### Neutralization Assay with SARS-COV-2 Spike-Pseudotyped Lentiviral Particles

For maximum antiviral effect, we design a combination containing 0.25 mg/mL Ai ye, 0.25 mg/mL, 0.25 mg/mL Jin lian hua, 0.25 mg/mL Jin yin hua and 0.25 mg/mL Hang bai ju. The anticoronavirus activity of the formula was further explored by pseudovirus neutralization test. Pseudoviruses containing the viral spike protein and expressing the luciferase reporter gene were used to infect embryonic kidney cells (HEK293) expressing human ACE2. The formula at the indicated concentration was incubated at room temperature for 90 minutes with spike glycoprotein pseudotyped viruses expressing luciferase expressed by the SARS-CoV-2 virus. Then, virus-containing the formula was incubated with ACE2-expressing HEK293 cells for 72 h, and viral infectivity was measured via luciferase. The incubation of pseudotyped lentivirus with the four kind volatile oil and the formula significantly reduced luciferase activities, and at 1 mg/ml of the highest inhibition was observed (Figure 4), confirming the ability of this formula to block entry of lentivirus spike protein into HEK293 cells either by direct inhibiting ACE2 or Cathepsin L or furin or Mpro/3cL protein on HEK293 cells. In addition, the effects of the formula on the cell viability were also evaluated, and it demonstrated no cytotoxicity (Figure 5)

**Fig 4.**
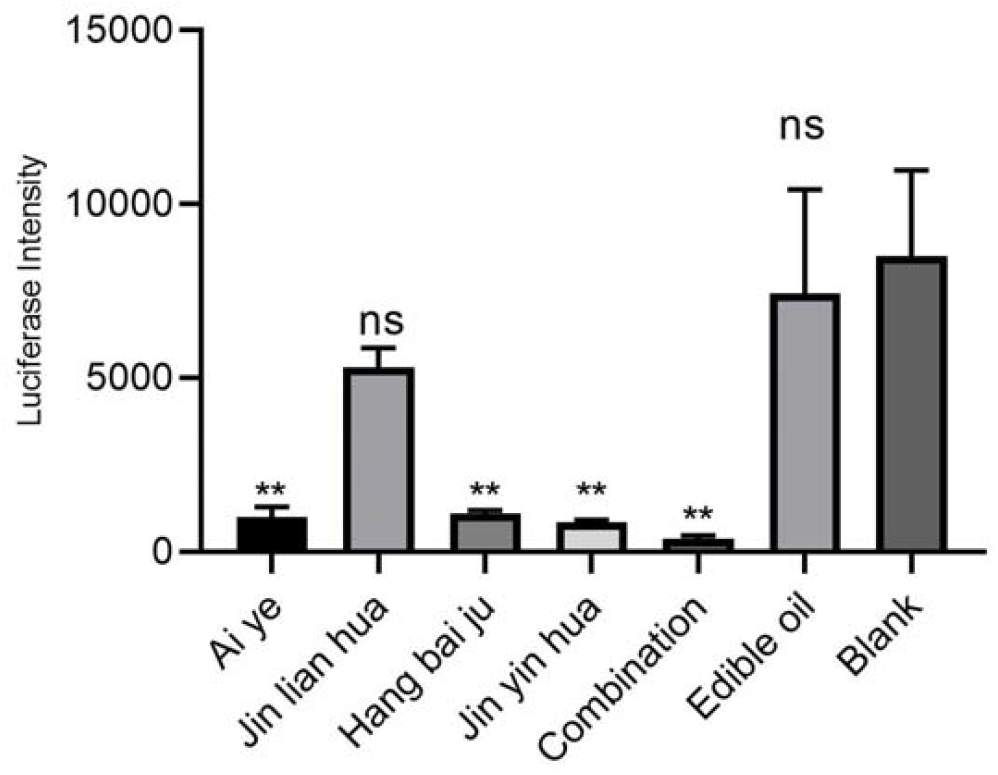
Combination of volatile oil inhibits SARS-CoV-2 pseudoviral infectivity in vitro.

**Fig 5.**
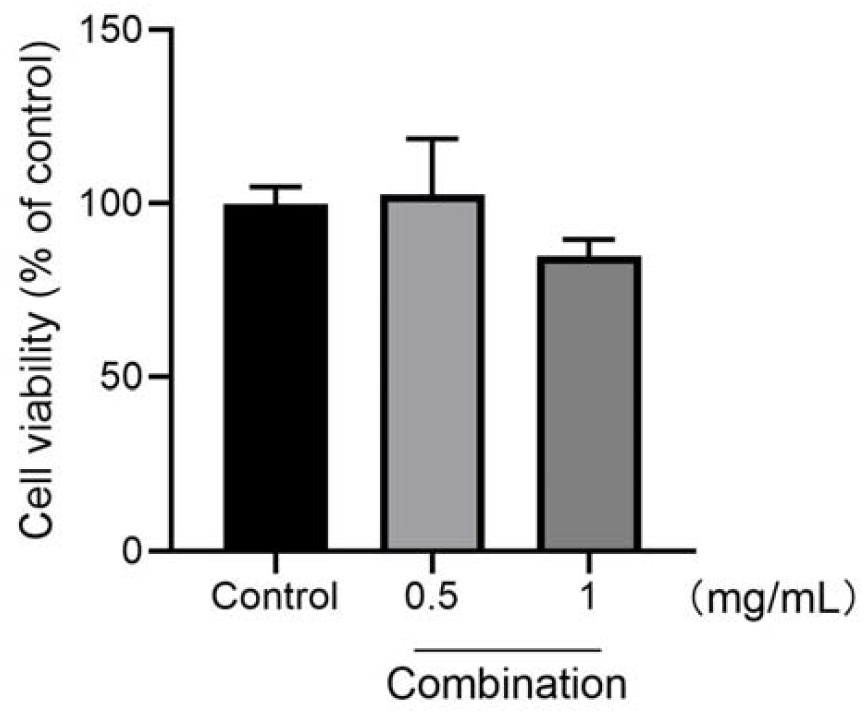
Cell viability were determined using CCK-8 assays. Values are percent changes above the control (DMSO, 0.1%) as the mean ± SD of three independent experiments. *p<0.05 and **p<0.01 compared to control cells.

The volatile oil in this article were rationally designed to target multiple targets either on viral entry or replication. The volatile oil in this article is a multiple target targeting COVID-19, This concept is consistent with the idea of drug discovery targeting COVID-19 multi protein reported previously^[15]^. The definition of a drug target is crucial to the success of drug discovery^[28]^. An antiviral agent targeting multiple viral proteins is a novel concept. Similar to drug cocktail screening, drug combination screening involves selecting drug candidates that target and block different stages of the virus’ life cycle. Contrary to the majority of existing repurposed drug research, this work focuses on multiple targets at once. Thus, our approach has the potential to attack the virus from different angles if successful.

### Exploring the Antiviral Mechanism of Formula Based on Molecular Docking

The antiviral mechanisms of four each volatile oil could preliminarily obtained through the literature. It is reported that the volatile oil of Ai ye has inhibitory effect on herpes zoster virus^[29]^, respiratory syncytial virus (RSV), influenza virus (IFV) and hepatitis B B virus (HBV)^[30,31]^. Artemisol and other components can inhibit bacteria and viruses^[31]^. Linalool, geraniol and a-terpineol from the volatile oil of Jin yin hua have antiviral effects^[32]^. Hai et al reported that volatile compounds are one of the main active ingredients of Hang bai ju, which have a significant anti influenza virus (H3N2) effect^[33,34]^. Zhao Hongwei et al. found that the ethanol extract of Jin lian hua has obvious inhibitory effect on virus replication in chicken embryos, and has direct killing effect on H1N1 virus PR8 strain^[35]^.

We next asked what is the specific mechanism underlying antiLJCOVID19 efficacy of the formula. It is necessary to explicitly determine the main components of these volatile oils. According to reports in the literature, volatile oil of Hang bai ju, Ai ye, Jin yin hua and Jin lian hua mainly contains 27^[36]^, 27^[37]^, 34^[38]^, 24^[39]^ components respectively, as shown in Table 2-5. To investigate the interactions between component of volatile oil and different targets, molecular docking experiments were performed. The component of the four kinds of essential oil were docked within the active site of four target using extra precision (XP) mode of Glide docking module of Maestro, respectively. Based on the docking scores, the compounds were ranked. PyMOL was used to visualize ligand-receptor interactions within the protein’s active site. The results shown that Isocyclocitral from Hang bai ju, spathulenol from Ai ye, benzyl benzoate from Jin yin hua, and beta-ionone from Jin lian hua bind at the active site of ACE2, furin, cathepainL and Mpro/3cL of SARS-CoV-2 with the dock score □-4.007, -4.995, -6.146 and □-4.927 kcal/mol, respectively (Table 2-5, Figure 6). Moreover, it also demonstrated that other components of each volatile oil could act on the corresponding target, even though the interaction between them is very weak.

**Table 2.** Molecular docking results of the essential Oil from Hang bai ju with ACE2.

| Number | ID | NAME | glide<br>ligand<br>efficiency | glide<br>gscore | glide<br>energy |
| --- | --- | --- | --- | --- | --- |
| 1 | CHEBI:171934 | Isocyclocitral | -0.364 | -4.007 | -17.951 |
| 2 | CHEBI:89773 | 1,2-Dihydro-1,1,6-trimethylnaphthalene | -0.277 | -3.596 | -17.596 |
| 3 | CHEBI:167345 | Cedrene | -0.238 | -3.564 | -19.948 |
| 4 | CHEBI:167347 | gamma-Selinene | -0.226 | -3.383 | -15.157 |
| 5 | CHEBI:10357 | (-)-beta-caryophyllene | -0.187 | -2.807 | -9.286 |
| 6 | CHEBI:180521 | D6-Ambrettolide | -0.154 | -2.763 | -12.213 |
| 7 | CHEBI:63190 | (+)-beta-caryophyllene | -0.179 | -2.68 | -10.828 |
| 8 | CHEBI:62224 | (-)-7-epi-alpha-selinene | -0.175 | -2.629 | -15.377 |
| 9 | CHEBI:62757 | alpha-curcumene | -0.169 | -2.529 | -17.887 |
| 10 | CHEBI:64361 | beta-sesquiphellandrene | -0.164 | -2.465 | -15.725 |
| 11 | CHEBI:84851 | ethyl linolenate | -0.085 | -1.862 | -30.722 |
| 12 | CHEBI:31572 | ethyl linoleate | -0.071 | -1.552 | -25.76 |
| 13 | CHEBI:34687 | dibutyl phthalate | -0.034 | -0.681 | -27.935 |
| 14 | CHEBI:32936 | tetracosane | 0.001 | 0.013 | -26.007 |
| 15 | CHEBI:46050 | docosane | 0.002 | 0.037 | -22.23 |
| 16 | CHEBI:17327 | phytol | 0.029 | 0.611 | -25.452 |
| 17 | CHEBI:39238 | (Z,E)-alpha-farnesene | 0.049 | 0.742 | -17.625 |
| 18 | CHEBI:17351 | linoleic acid | 0.071 | 1.42 | -22.532 |
| 19 | CHEBI:69080 | methyl linoleate | 0.072 | 1.515 | -25.416 |
| 20 | CHEBI:10418 | trans-beta-farnesene | 0.111 | 1.669 | -18.428 |
| 21 | CHEBI:30813 | decanoic acid | 0.161 | 1.927 | -14.954 |
| 22 | CHEBI:15756 | hexadecanoic acid | 0.133 | 2.394 | -21.115 |
| 23 | CHEBI:43619 | icosane | 0.154 | 3.077 | -22.779 |
| 24 | CHEBI:16148 | heptadecane | 0.189 | 3.221 | -20.402 |
| 25 | CHEBI:32927 | nonadecane | 0.171 | 3.249 | -21.935 |
| 26 | CHEBI:32926 | octadecane | 0.189 | 3.405 | -20.733 |
| 27 | CHEBI:132293 | 2-methylicosane | 0.162 | 3.407 | -21.296 |

**Table 3.**
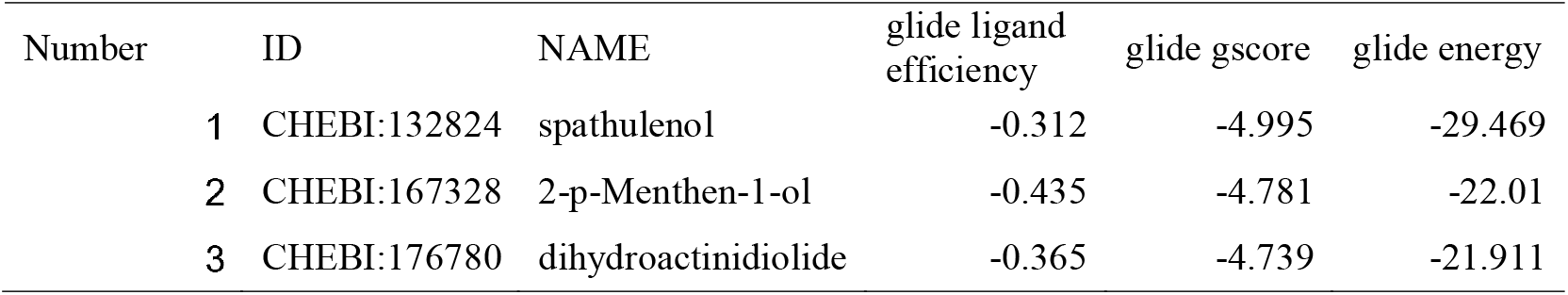
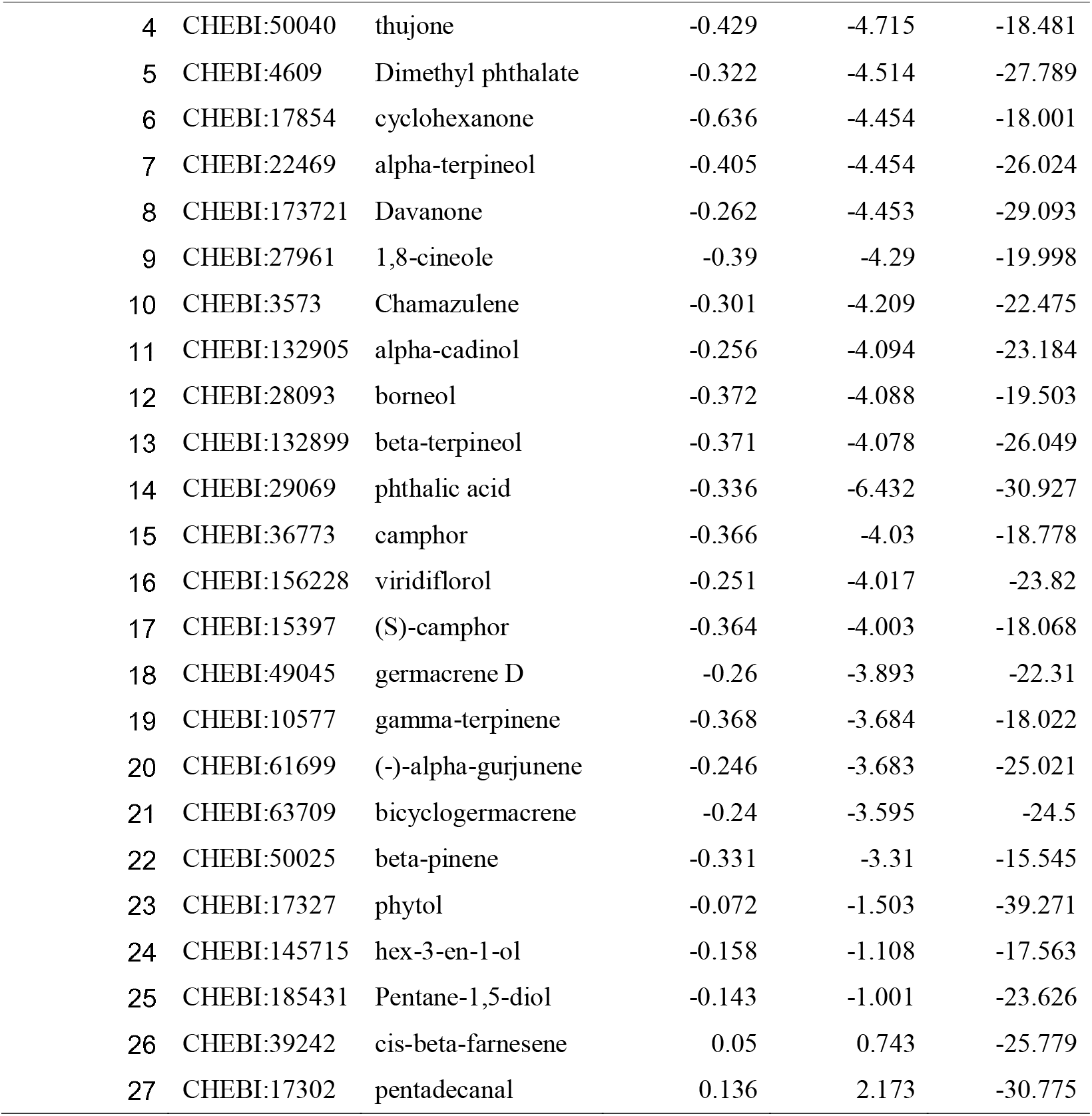
Molecular docking results of the essential Oil from Ai ye with furin.

| Number | ID | NAME | glide ligand<br>efficiency | glide gscore | glide energy |
| --- | --- | --- | --- | --- | --- |
| 1 | CHEBI:132824 | spathulenol | -0.312 | -4.995 | -29.469 |
| 2 | CHEBI:167328 | 2-p-Menthen-1-ol | -0.435 | -4.781 | -22.01 |
| 3 | CHEBI:176780 | dihydroactinidiolide | -0.365 | -4.739 | -21.911 |
| 4 | CHEBI:50040 | thujone | -0.429 | -4.715 | -18.481 |
| 5 | CHEBI:4609 | Dimethyl phthalate | -0.322 | -4.514 | -27.789 |
| 6 | CHEBI:17854 | cyclohexanone | -0.636 | -4.454 | -18.001 |
| 7 | CHEBI:22469 | alpha-terpineol | -0.405 | -4.454 | -26.024 |
| 8 | CHEBI:173721 | Davanone | -0.262 | -4.453 | -29.093 |
| 9 | CHEBI:27961 | 1,8-cineole | -0.39 | -4.29 | -19.998 |
| 10 | CHEBI:3573 | Chamazulene | -0.301 | -4.209 | -22.475 |
| 11 | CHEBI:132905 | alpha-cadinol | -0.256 | -4.094 | -23.184 |
| 12 | CHEBI:28093 | borneol | -0.372 | -4.088 | -19.503 |
| 13 | CHEBI:132899 | beta-terpineol | -0.371 | -4.078 | -26.049 |
| 14 | CHEBI:29069 | phthalic acid | -0.336 | -6.432 | -30.927 |
| 15 | CHEBI:36773 | camphor | -0.366 | -4.03 | -18.778 |
| 16 | CHEBI:156228 | viridiflorol | -0.251 | -4.017 | -23.82 |
| 17 | CHEBI:15397 | (S)-camphor | -0.364 | -4.003 | -18.068 |
| 18 | CHEBI:49045 | germacrene D | -0.26 | -3.893 | -22.31 |
| 19 | CHEBI:10577 | gamma-terpinene | -0.368 | -3.684 | -18.022 |
| 20 | CHEBI:61699 | (-)-alpha-gurjunene | -0.246 | -3.683 | -25.021 |
| 21 | CHEBI:63709 | bicyclogermacrene | -0.24 | -3.595 | -24.5 |
| 22 | CHEBI:50025 | beta-pinene | -0.331 | -3.31 | -15.545 |
| 23 | CHEBI:17327 | phytol | -0.072 | -1.503 | -39.271 |
| 24 | CHEBI:145715 | hex-3-en-1-ol | -0.158 | -1.108 | -17.563 |
| 25 | CHEBI:185431 | Pentane-1,5-diol | -0.143 | -1.001 | -23.626 |
| 26 | CHEBI:39242 | cis-beta-farnesene | 0.05 | 0.743 | -25.779 |
| 27 | CHEBI:17302 | pentadecanal | 0.136 | 2.173 | -30.775 |

**Table 4.** Molecular docking results of the essential Oil from Jin yin hua with Cathepsin L.

| Number | ID | NAME | glide<br>ligand<br>efficiency | glide<br>gscore | glide<br>energy |
| --- | --- | --- | --- | --- | --- |
| 1 | CHEBI:41237 | benzyl benzoate | -0.384 | -6.146 | -32.778 |
| 2 | CHEBI:28266 | fluorene | -0.454 | -5.901 | -24.18 |
| 3 | CHEBI:22469 | alpha-terpineol | -0.535 | -5.89 | -25.072 |
| 4 | CHEBI:28851 | phenanthrene | -0.416 | -5.83 | -24.734 |
| 5 | CHEBI:34247 | 2,6-di-tert-butyl-4-methylphenol | -0.342 | -5.472 | -23.883 |
| 6 | CHEBI:32319 | alpha-ionone | -0.35 | -4.906 | -23.664 |
| 7 | CHEBI:84851 | ethyl linolenate | -0.17 | -3.745 | -36.322 |
| 8 | CHEBI:32936 | tetracosane | -0.15 | -3.595 | -35.184 |
| 9 | CHEBI:17447 | geraniol | -0.325 | -3.575 | -23.114 |
| 10 | CHEBI:32941 | heptacosane | -0.114 | -3.082 | -39.933 |
| 11 | CHEBI:32938 | pentacosane | -0.106 | -2.659 | -34.403 |
| 12 | CHEBI:17580 | linalool | -0.236 | -2.598 | -21.637 |
| 13 | CHEBI:149547 | (2E,4E)-deca-2,4-dienal | -0.233 | -2.568 | -24.047 |
| 14 | CHEBI:46050 | docosane | -0.1 | -2.211 | -35.118 |
| 15 | CHEBI:34687 | dibutyl phthalate | -0.098 | -1.956 | -39.578 |
| 16 | CHEBI:67252 | farnesyl acetone | -0.097 | -1.85 | -32.469 |
| 17 | CHEBI:17351 | linoleic acid | -0.092 | -1.846 | -38.83 |
| 18 | CHEBI:177721 | Methyl 9-oxononanoate | -0.073 | -0.946 | -29.415 |
| 19 | CHEBI:17327 | phytol | -0.033 | -0.683 | -32.077 |
| 20 | CHEBI:42504 | pentadecanoic acid | -0.039 | -0.664 | -33.772 |
| 21 | CHEBI:15756 | hexadecanoic acid | -0.024 | -0.442 | -36 |
| 22 | CHEBI:28842 | octadecanoic acid | -0.02 | -0.397 | -39.091 |
| 23 | CHEBI:30805 | dodecanoic acid | -0.028 | -0.396 | -29.905 |
| 24 | CHEBI:30813 | decanoic acid | -0.033 | -0.394 | -28.055 |
| 25 | CHEBI:69187 | Methyl palmitate | -0.015 | -0.277 | -33.707 |
| 26 | CHEBI:69080 | methyl linoleate | -0.011 | -0.224 | -35.128 |
| 27 | CHEBI:28875 | tetradecanoic acid | 0 | -0.004 | -30.723 |
| 28 | CHEBI:69188 | Methyl stearate | 0.006 | 0.136 | -36.556 |
| 29 | CHEBI:84156 | methyl palmitoleate | 0.016 | 0.312 | -33.112 |
| 30 | CHEBI:84067 | tetradecanal | 0.064 | 0.956 | -29.986 |
| 31 | CHEBI:45296 | hexadecane | 0.111 | 1.774 | -28.734 |
| 32 | CHEBI:32927 | nonadecane | 0.099 | 1.884 | -30.142 |

**Table 5.**
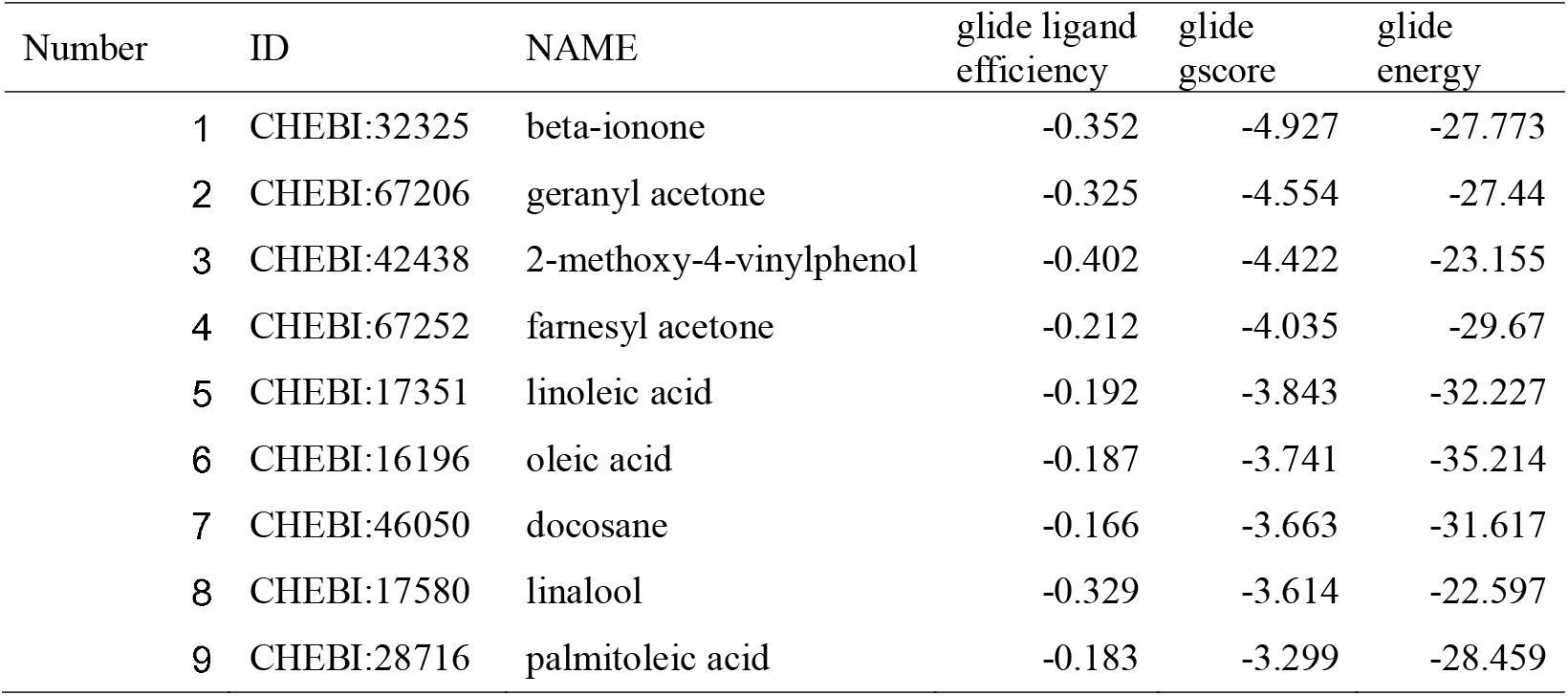
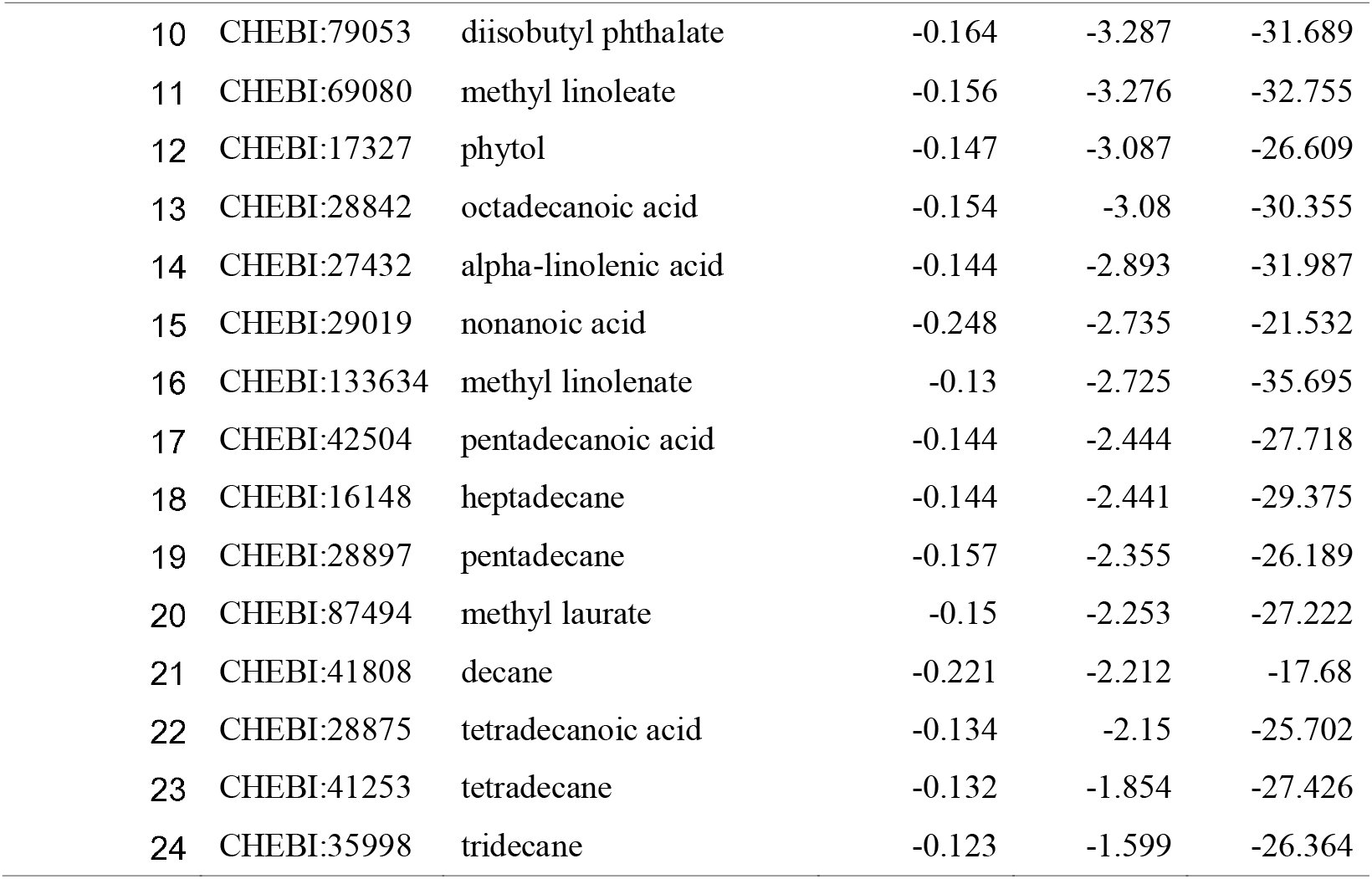
Molecular docking results of the essential Oil from Jin lian hua with Mpro/3cl.

**Fig 6.**
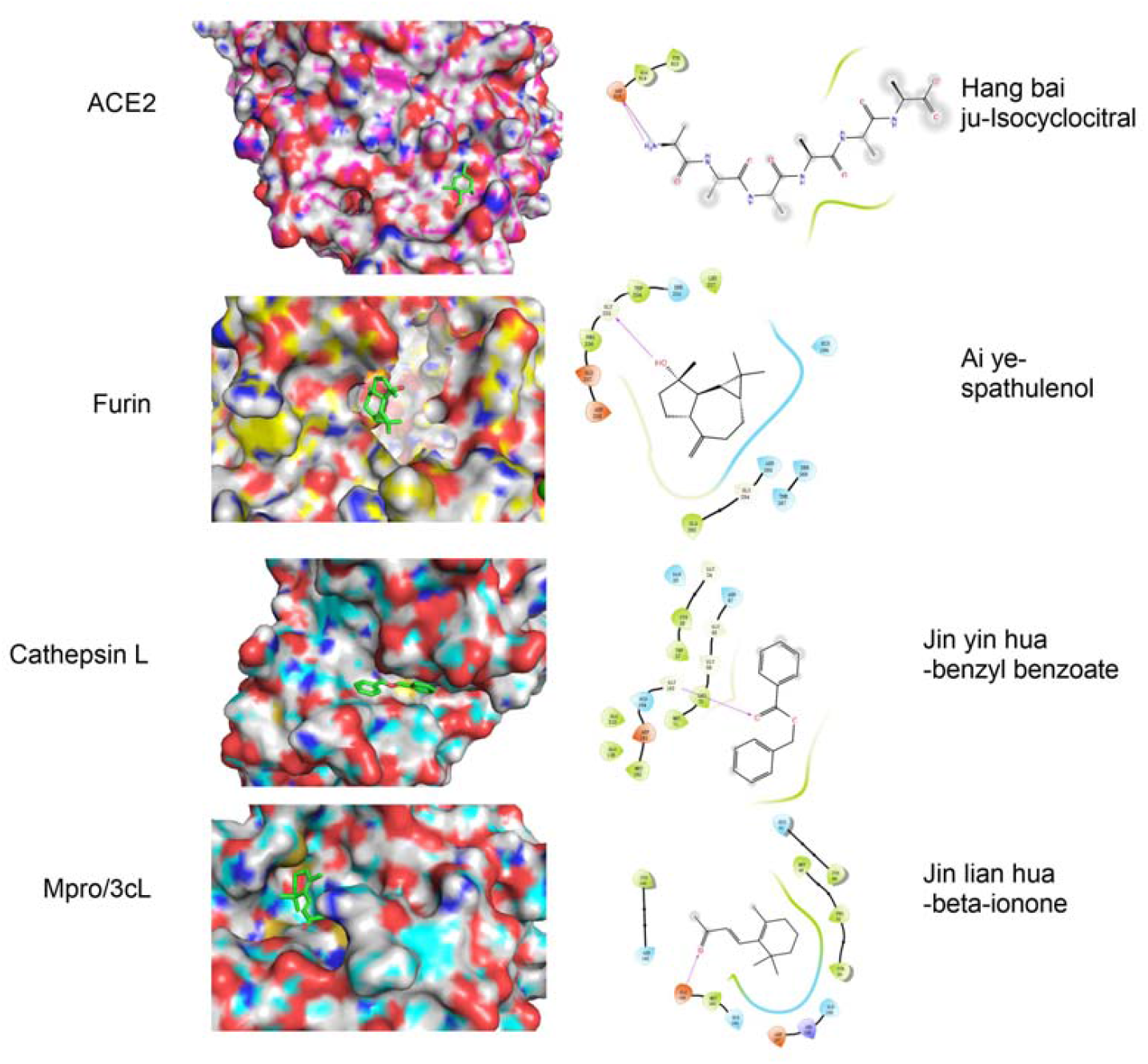
Molecular docking of representative components of Jin yin hua, Jin lian hua, Ai ye and Hang bai ju with Cathespin L, Mpro/3cL, Furin and ACE2 Proteins, respectively

There are only five components of the volatile oil from Jin yin hua, whose docking score with absolute values were greater than 5, indicating that the binding ability of these volatile oil monomers to the target was weak. As is well known to all, herbs produce their efficacy through the synergistic effects of multi-ingredients, multi-targets, and multi-pathways^[40,41]^. Although the results show that the absolute value of the docking score between these volatile oil monomers and the target is relatively small, these volatile oils with multi-component synergizing on the common target can also achieve the effect of acting on the target.

## Conclusions

In this work, we present a formula consisting volatile oil for anti-viral agents that interact with multiple target proteins of SARS-CoV-2. We first collected 9 antiviral volatile oils from the literature, Secondly, the volatile oil with the best inhibitory effect on the four targets was obtained by target screening. Then the formula consisting the four volatile oils has better effect of neutralizing pseudovirus compared with the effects of mono-agents. Overall, we found a volatile oil formula with potential inhibitory ability against 4 key proteins of SARS-CoV-2. The volatile oil can be used to develop oral or nasal preparations. A further study in animals is needed to determine if oral or nasal spray delivery is more effective as a preventative or curative measure. There are more than four targets related to the life cycle of novel coronavirus^[6]^. In the future, we will further screen other targets to improve the efficacy of this formula.

## Author Contribution

Conceptualization: HZ and BZ; methodology and resources: HZ, WG, JS, and XS; validation and visualization: HZ and WG; formal analysis: HZ, WG and BZ; writing—original draft preparation: HZ; writing—review and editing: HZ, WG and BZ; supervision: BZ. All authors have read and agreed to the published version of the manuscript.

## Data Availability

The data used to support the fndings of this study are available from the corresponding author upon request.

## Declarations

### Ethics Approval

Not applicable.

### Consent to Participate

Not applicable.

### Consent for Publication

Not applicable.

### Confict of Interest

The authors declare no competing interests.

### Open Access

This article is licensed under a Creative Commons Attribution 4.0 International License, which permits use, sharing, adaptation, distribution and reproduction in any medium or format, as long as you give appropriate credit to the original author(s) and the source, provide a link to the Creative Commons licence, and indicate if changes were made. The images or other third party material in this article are included in the article’s Creative Commons licence, unless indicated otherwise in a credit line to the material. If material is not included in the article’s Creative Commons licence and your intended use is not permitted by statutory regulation or exceeds the permitted use, you will need to obtain permission directly from the copyright holder. To view a copy of this licence, visit http://creativecommons.org/licenses/by/4.0/.

